# A clinically validated human capillary blood transcriptome test for global systems biology studies

**DOI:** 10.1101/2020.05.22.110080

**Authors:** Ryan Toma, Ben Pelle, Nathan Duval, Matthew M Parks, Vishakh Gopu, Hal Tily, Andrew Hatch, Ally Perlina, Guruduth Banavar, Momchilo Vuyisich

## Abstract

Chronic diseases are the leading cause of morbidity and mortality globally. Yet, the majority of them have unknown etiologies, and genetic contribution is weak. In addition, many of the chronic diseases go through the cycles of relapse and remission, during which the genomic DNA does not change. This strongly suggests that human gene expression is the main driver of chronic disease onset and relapses. To identify the etiology of chronic diseases and develop more effective preventative measures, a comprehensive gene expression analysis of the human body is needed. Blood tissue is easy to access and contains a large number of expressed genes involved in many fundamental aspects of our physiology.

We report here the development of a whole blood transcriptome clinical test that is high throughput, automated, inexpensive, and clinically validated. The test requires only 50 microliters of blood from a finger prick, enabling access by diverse populations that have been traditionally under-represented in clinical research. The transcripts in the samples are preserved at the time of collection and can be stored and/or transported at ambient temperatures for up to 28 days. The sample preservative protects integrity, while also inactivating all pathogens (bacteria, fungi, and viruses), enabling safe transportation globally. Given its unique set of usability features and clinical performance, this test should be integrated into longitudinal, population-scale, systems biology studies.

## Introduction

Quantitative gene expression analysis provides a global snapshot of tissue function. The human transcriptome varies with tissue type, developmental stage, environmental stimuli, and health/disease state (Frith et al., 2005; Lin et al., 2019; Marioni et al., 2008; Mortazavi et al., 2008; Velculescu et al., 1999). Changes in the expression patterns can provide insights into the molecular mechanisms of disease onset and progression. In the era of precision medicine, methods to non-invasively assess an individual’s transcriptional changes can reveal predictive markers of disease, inform the best choice of therapy, and enable the development of novel therapies. For example, using systems biology approaches to interrogate the transcriptome are now a standard strategy to stratify cancer patients and select the best available therapies (Cabanero and Tsao, 2018; Chauhan et al., 2019; Rosell et al., 2012). Methods to analyze the transcriptome have become a vital part of trying to better understand chronic diseases with unknown etiologies.

In the United States, chronic, non-communicable diseases have become an epidemic with an ever-growing share of health care expenditures (CDC, 2020). Many chronic diseases go through cycles of elevated disease activity (disease relapse) and periods of low disease activity (disease remission) (Carlberg et al., 2019; Haberman et al., 2019; Ostrowski et al., 2019; Stephenson et al., 2018; Zhao et al., 2013). The human genome has provided valuable information on disease predisposition and susceptibility, but the overall genetic contribution to many chronic diseases is weak (Rappaport, 2016). Furthermore, many environmental or lifestyle factors increase the risk of developing a chronic disease, implicating a change in gene expression. This has been shown in large twin and epidemiological studies (Hand et al., 2016; Wilkins et al., 2019). Changes in gene expression have been shown in type I diabetes (Reynier et al., 2010), type II diabetes (Berisha et al., 2011; Homuth et al., 2015), and cardiovascular disease (Pedrotty et al., 2012; Wirka et al., 2018), often associated with immune regulation (Chaussabel et al., 2010). Unfortunately, little progress has been made in understanding the role of the transcriptome in chronic disease due to the lack of a low-cost and scalable method for which samples can be collected by anyone and anywhere, instead relying on complex logistics of collecting peripheral blood at a clinic and transporting it frozen to a laboratory. In addition, there are very few clinically validated blood transcriptome tests available, so analyses of data from disparate methods can lead to erroneous conclusions due to biases introduced in each method.

Circulating blood is easily accessible and provides a non-invasive alternative to tissue biopsies for molecular profiling of human disease and disease risk (Liew et al., 2006). Peripheral blood mononuclear cells (PBMC) have been the primary choice for transcriptomic investigations from whole blood samples (Sen et al., 2018). Unfortunately, methods available for scrutinizing the transcriptome from PBMCs are useful for research methods, but not ideal for broader applications at scale. Furthermore, methods using PBMCs fail to measure the cell-free transcripts, which have been associated with certain disease conditions, particularly for cancer patients and pregnant women (Ng et al., 2003; Pinzani et al., 2010; Pös et al., 2018; Souza et al., 2017).

The current state of the art focuses on the whole blood transcriptome, typically requiring a venous blood draw. The availability of high-throughput sequencing technologies has enabled the rapid analysis of the transcriptome from small sample volumes. To our knowledge, there is currently no easily scalable blood test for population-based studies using whole-blood transcriptome analysis. Here we describe, for the first time, a high-throughput method for interrogating the human transcriptome from a small finger prick blood sample. The method is automated, inexpensive, and clinically validated. An RNA preservation buffer mixed with the blood sample at the point of collection inactivates all pathogens (bacteria, fungi, and viruses), enabling safe transportation globally at ambient temperatures for up to 28 days, while preserving RNA integrity for gene expression analysis.

## Methods

### Ethics statement

All procedures involving human participants were performed in accordance with the ethical standards and approved by a federally accredited Institutional Review Board (IRB) committee. Informed consent was obtained from all participants.

### Sample collection

Capillary blood was collected by study participants using an at-home kit, following the included instructions. The kit contains a 1.5mL screw cap tube (Sarstedt) containing 200uL of RNA preservative buffer (RPB), a 50uL EDTA-coated minivette (Sarstedt), a 70% isopropyl alcohol wipe (Dynarex), a gauze pad, a push-button 17-gauge disposable lancet (Acti-Lance), and instruction sheet. Participants were instructed to fast for 8 hours prior to collection and collect the samples within one hour of waking up, without exercising or drinking (except water). The finger was wiped with a 70% isopropyl alcohol wipe and allowed to dry. Using the disposable lancet, the finger was punctured, and the first drop wiped away using the gauze pad. The blood was drawn by capillary action into a 50uL EDTA-coated minivette and dispensed into the sample collection tube that contained 200uL of RPB. The tube was vigorously shaken for 15 seconds to thoroughly mix RPB with blood. The puncture was wiped clean and a band-aid was applied.

Peripheral (venous) blood was collected by a phlebotomist. Participants were instructed to fast for 8 hours prior to collection and collect the samples within one hour of waking up, without exercising or drinking (except water). The puncture site was first cleaned with a 70% isopropyl alcohol wipe (Dynarex) and allowed to dry. Blood was then collected into 4 mL EDTA vacutainers (BD Bioscience). The blood was promptly transferred from the vacutainers to another tube and mixed with RPB at a 1:4 ratio.

Samples used to compare gene expression in peripheral and capillary blood were collected using the methods described above. For six participants, peripheral and capillary blood samples were collected within minutes of each other.

Longitudinal samples were collected weekly from 10 participants over a five-week period using the capillary blood collection method described above. Four technical replicates were collected and analyzed at each time point.

Shipping, storage, and precision validation samples were collected from three participants following the capillary blood collection method described above. To validate the storage stability, samples were stored for 7, 14, and 28 days at room temperature (RT) or −80°C and compared to the technical replicates analyzed the day of collection (day 0). A subset of replicate samples was shipped twice during storage times.

### RNA extraction, library preparation, and sequencing

RNA was extracted using silica beads and a series of washes, followed by elution in molecular biology grade water. DNA was degraded using RNase-free DNase. Polyadenylated messenger RNAs (mRNAs) were isolated from the total RNA using an oligo dT-based magnetic mRNA isolation kit. mRNAs were converted to directional sequencing libraries with unique dual-barcoded adapters and ultrapure reagents. Libraries were pooled and quality controlled with dsDNA Qubit (Thermo Fisher Scientific), QuantStudio 3 qPCR (Thermofisher) with KAPA Library Quantification Kit (Roche), and Fragment Analyzer (Advanced Analytical). Library pools were sequenced on Illumina NextSeq or NovaSeq instruments using 300 cycle kits.

### Bioinformatics

A human transcriptome catalog was created by augmenting the transcripts derived from the Ensembl release 99 (Yates et al., 2019) annotations for GRCh38 (Schneider et al., 2017) with selected transcripts from RefSeq. Raw reads were mapped to this transcript catalog and quantified with Salmon (Patro et al., 2017). Transcript expression was aggregated by gene to obtain gene expression levels. In a sample, only genes with a read count of at least one were considered as expressed. Transcripts per million for genes were calculated using the expected read counts and effective read lengths as generated by the statistical model in Salmon.

### Data analysis

Statistical parameters, including transformations and significance, are reported in the figures and figure legends. To compare pairs of samples, we report Jaccard similarity (which ignores expression and considers overlap in genes detected), Spearman correlations (which are invariant to absolute expression levels of the genes and only consider the similarity of ranked expression), Pearson correlations on logged data (which measure the linear relationship between gene expression levels), and Hellinger distance (an appropriate distance measure for compositional data). Pearson and Spearman correlation coefficients were performed on centered log ratio (CLR) transformed data (Aitchison, 1982), as is commonly done to reduce false discoveries due to the compositional nature of sequencing data. Statistical analysis was performed in python.

## Results

### Correlation of gene expression levels in capillary and peripheral blood

Peripheral (venous) blood is the current gold standard sample for whole blood transcriptome analysis, so we compared the gene expression levels obtained from peripheral and capillary blood of six study participants. The correlation between gene expression in capillary and peripheral blood from these participants shows excellent concordance, by Spearman correlation coefficient on center log ratio (CLR) transformed values above 0.94 for all donors (Figure 2). Transcripts from an average of nearly 14,000 genes are detected in each sample.

**Figure 1.**
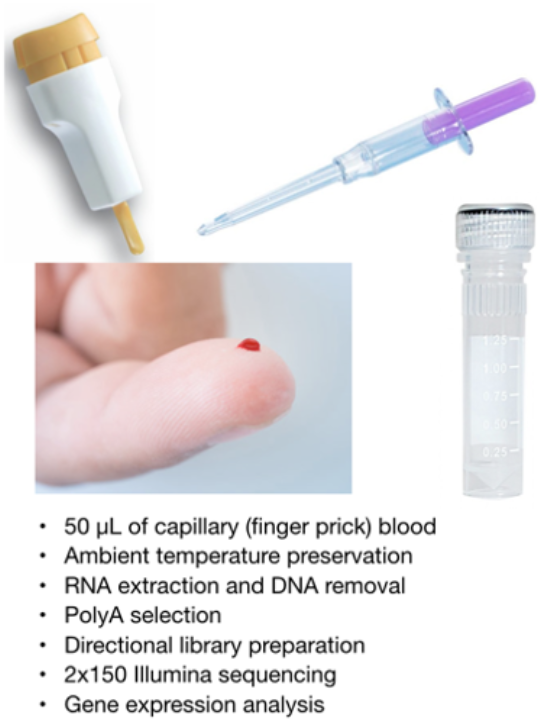
Primary components of the at-home capillary blood collection kit and outline of the sample analysis method. The lancet is used to prick the finger; the minivette is a capillary device with a piston; the sample collection tube is a microcentrifuge tube with 200 microliters of RPB.

**Figure 2.**
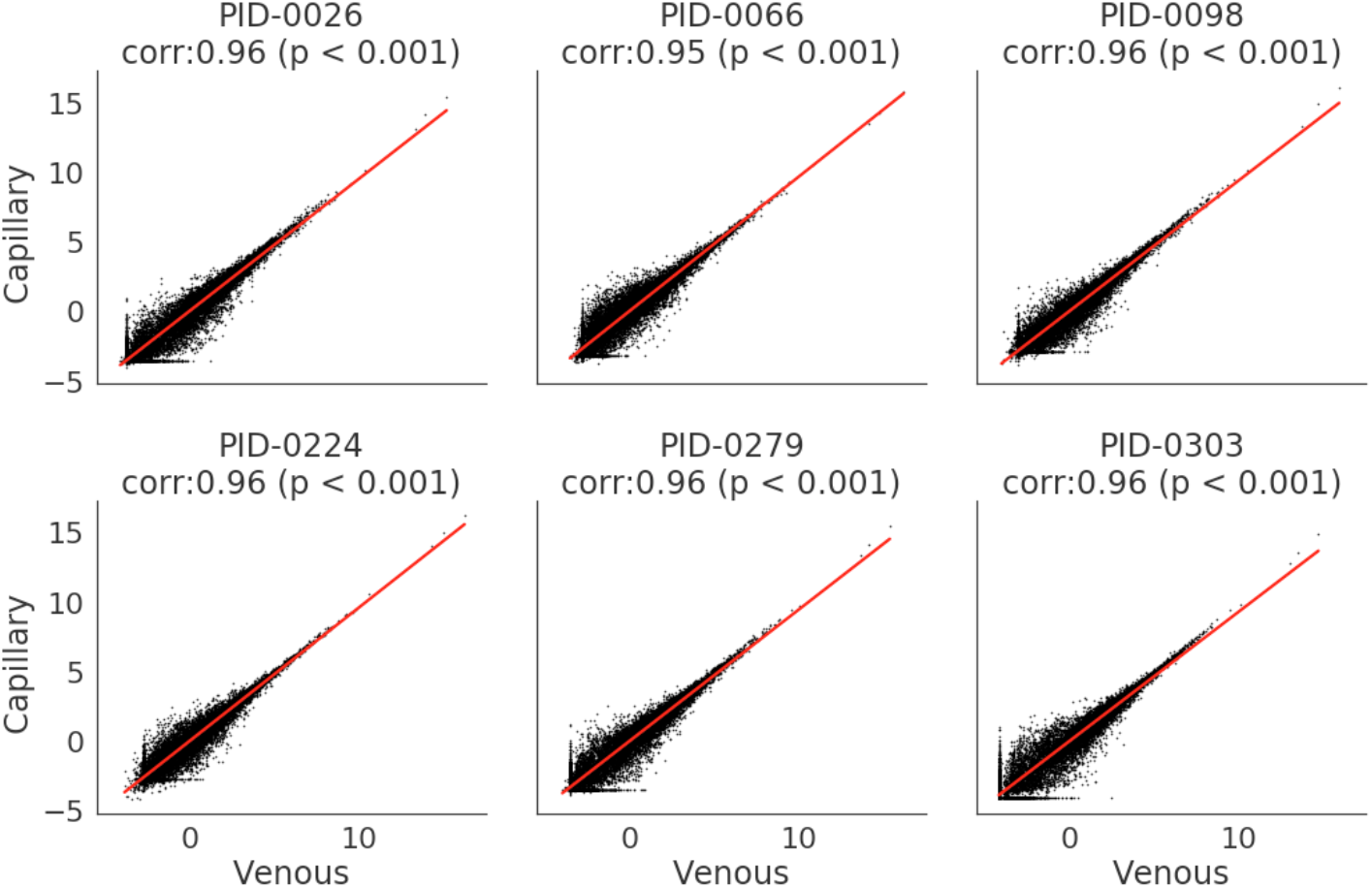
Correlation between peripheral and capillary blood gene expression levels from six donors (PID-####) shows high concordance. Spearman correlation coefficient on center log ratio (CLR) transformed values are above 0.94 for all instances with p-values < 0.001.

### Method precision

For clinical studies in a large population and over time that measure gene expression changes as a function of health and disease states, method precision is a very important parameter. To determine the precision of the method, we measured the percent coefficient of variation (%CV) for gene expression in four technical replicates from three participants (Figure 3). The %CV is plotted as a function of expression level (mean counts per million, CPM), and as expected, the higher the expression level, the lower the variance. The distribution of transcript relative abundance is shown from 10 CPM to greater than 10^5^ CPM. A small cluster greater than 10^5^ CPM are highly abundant globin transcripts. A CV below 25% for the majority of transcripts shows low variance for this method.

**Figure 3.**
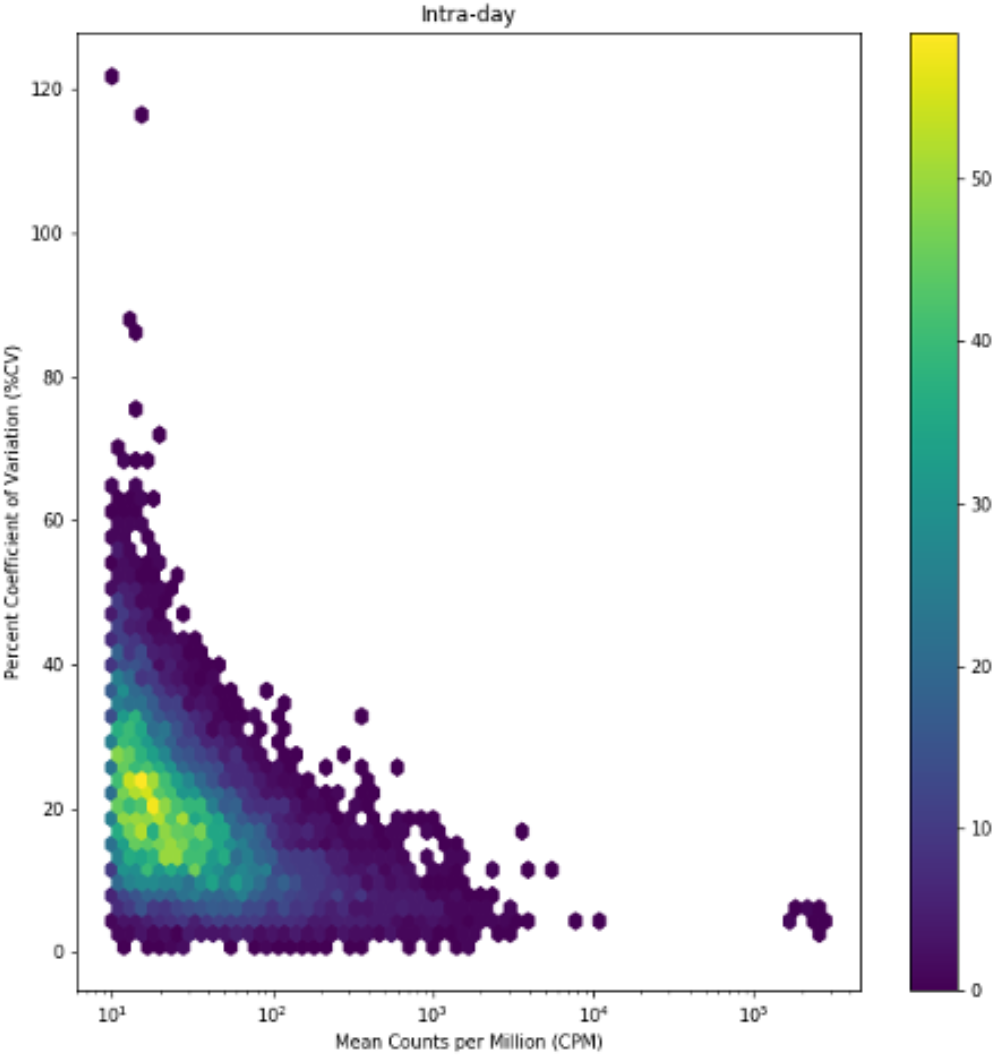
The method precision is shown as the percent coefficient of variation (%CV) as a function of transcript abundance (mean CPM). These data show a broad distribution of transcript abundances, from 10 CPM to greater than 10^5^ CPM.

### Sample Stability

One concern when performing transcriptomic analysis of blood samples is the facile degradation of RNA, due to self-cleavage and action of ribonucleases (Souza et al., 2017). To preserve RNA, our method uses an RNA preservative buffer (RPB) with multiple functions. RPB’s detergents immediately dissolve lipid bilayers and inactivate all pathogens, the denaturants halt all enzymatic activity by disrupting the tertiary structures of proteins, and special buffering agents prevent RNA self-cleavage by keeping the 2’ oxygen protonated, thus preserving the RNA integrity during sample shipping and storage. To validate the efficacy of RPB to preserve the blood transcriptome at ambient temperatures, blood samples were collected from three participants and stored at ambient temperature (72°F, 22°C) or −80°C for 0, 7, 14, and 28 days, with and without shipping. Four technical replicates per condition were analyzed using our test and gene expression levels compared among all samples. The correlation between blood samples in all conditions is very high, with Spearman correlation coefficients above 0.81 for all conditions tested (Akoglu, 2018). Among all comparisons, the lowest spearman correlation for donor 1 was 0.849 (Figure 4), the lowest spearman correlation for donor 2 was 0.865 (Figure S1), and the lowest spearman correlation for donor 3 was 0.811 (Figure S1). This shows that our method can adequately preserve RNA for up to 28 days at ambient temperature.

**Figure 4.**
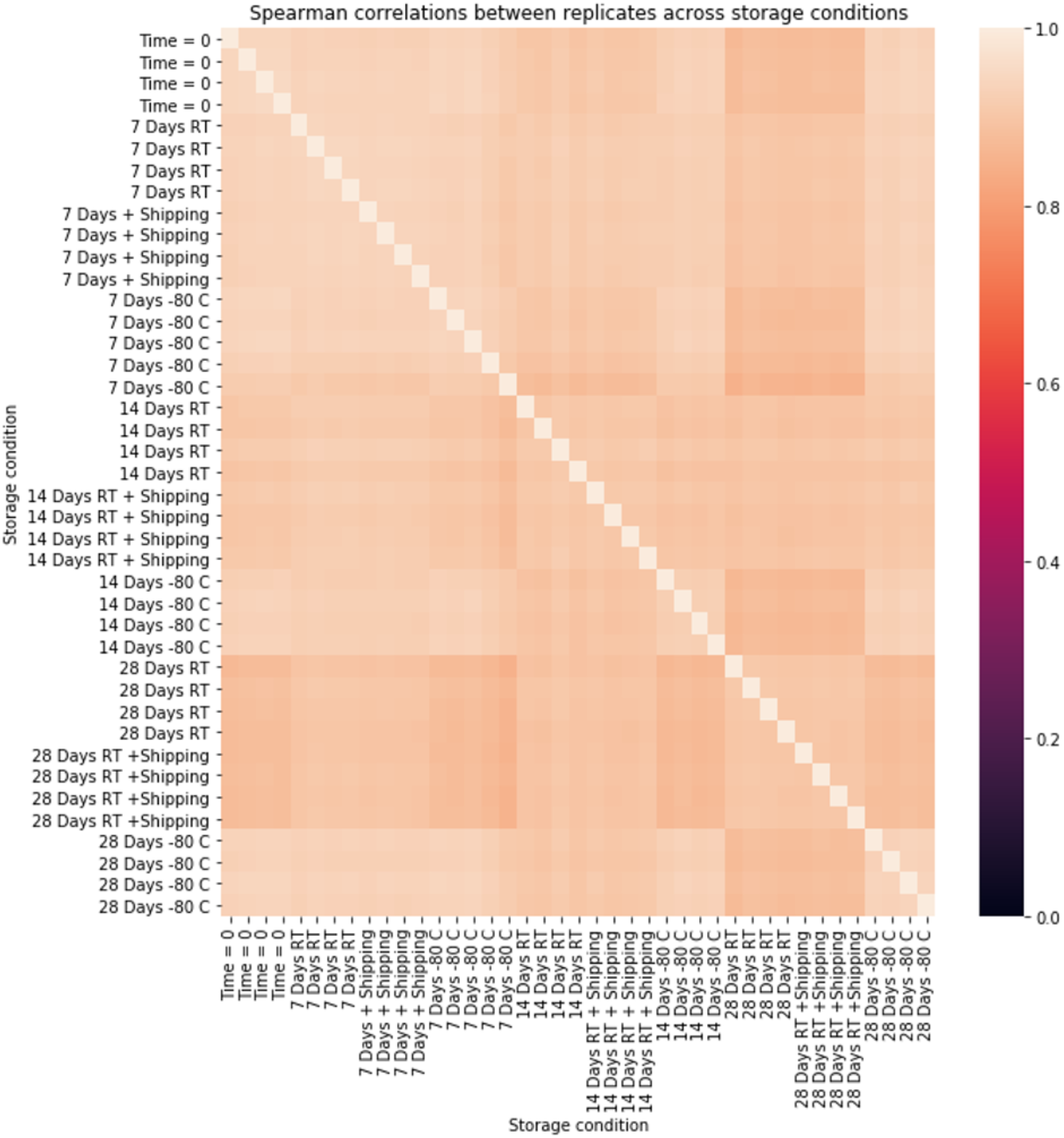
Heatmap showing Spearman correlation coefficients for sample stability of blood samples stored at ambient temperature and −80°C for 0, 7, 14, and 28 days, with and without shipping conditions. The Spearman correlation coefficients for all conditions in all donor samples were greater than 0.81.

### Longitudinal changes in the blood transcriptome

In this small study, 10 participants collected capillary blood weekly for five weeks. On average, transcripts from 13,796 genes were detected per sample. For genes that are detected, the correlation between gene expression remains high across five weeks for each participant. Jaccard similarity, which measures the overlap in genes detected at all between samples, is stable at around 50%. This suggests that while around half the genes detected in any sample may not be detected in another sample from the same donor (these are close to the limit of detection), the genes that are consistently detected are expressed at a relatively stable level (Figure 5).

**Figure 5.**
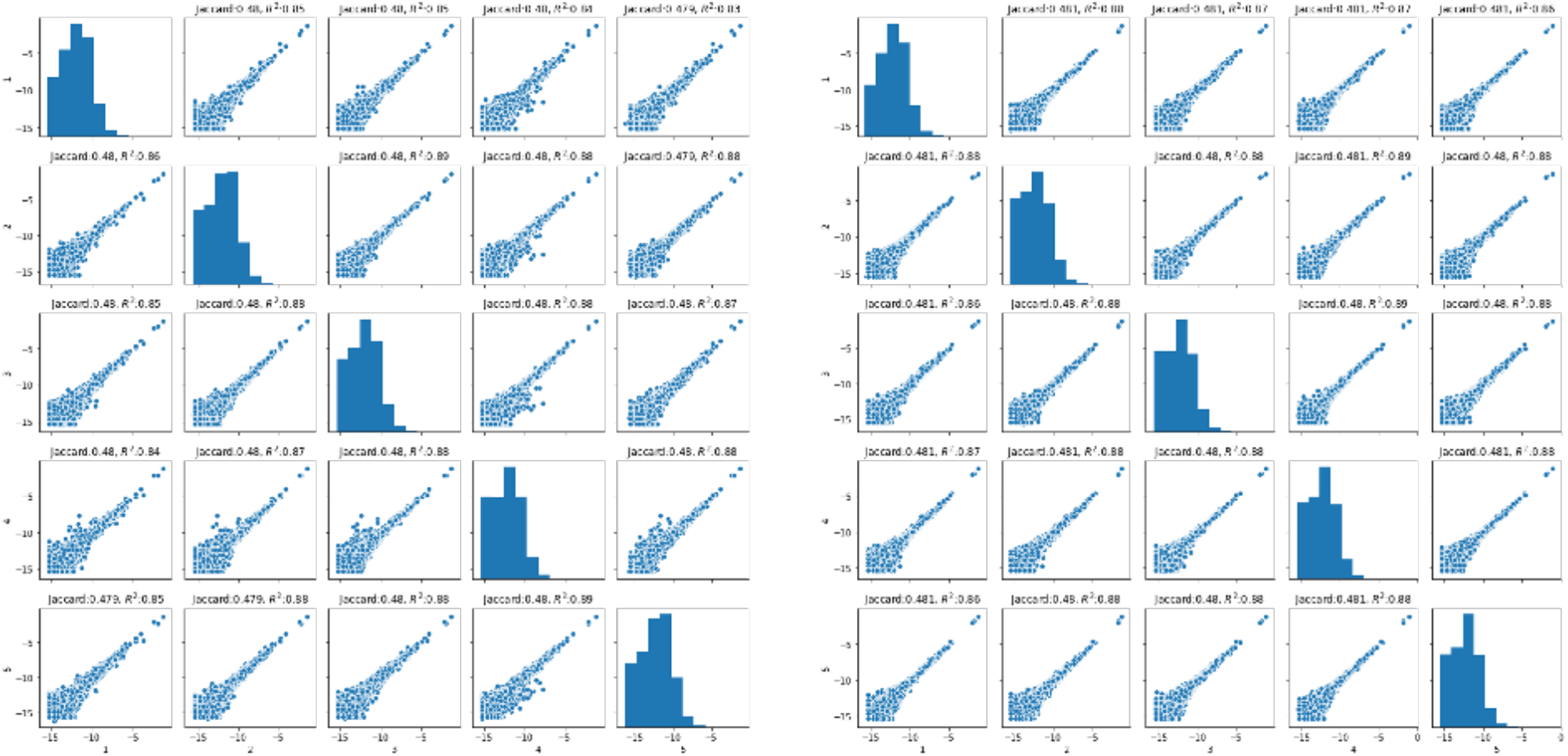
Longitudinal stability of whole blood transcriptome in two study participants over a 5-week period. Pearson correlation and Jaccard similarity between pairs of samples, one sample randomly selected each week over five weeks, for two randomly chosen participants. Correlations are calculated on log-transformed values and only for genes that are detected in both samples.

One of the more important parameters of the blood test is to be sufficiently precise to distinguish very small changes among thousands of measured features. Such high precision would enable the identification of transcriptome changes over time and due to changes in health and disease. To assess the test precision in this context, we compared Hellinger distances among 1. Technical replicates from samples obtained on the same day from the same person (intraperson technical distances in Figure 6), 2. Biological samples from the same person collected over time (intraperson distances in Figure 6), and 3. Biological samples obtained from different people (interperson distances in Figure 6). Empirical cumulative distribution function (ECDF) plot of Hellinger distances shows that when taking all genes into account, pairs of samples taken from an individual within one week (intra-person - technical, green) are more similar to samples from that same individual taken on two different weeks (intra-person, orange). However, pairs of samples coming from two different participants (inter-person, blue) tend to be less similar than samples taken from a single participant (Figure 6). Furthermore, the Hellinger distances show that although less similar, there are still a substantial number of genes that are stably expressed between individuals.

**Figure 6.**
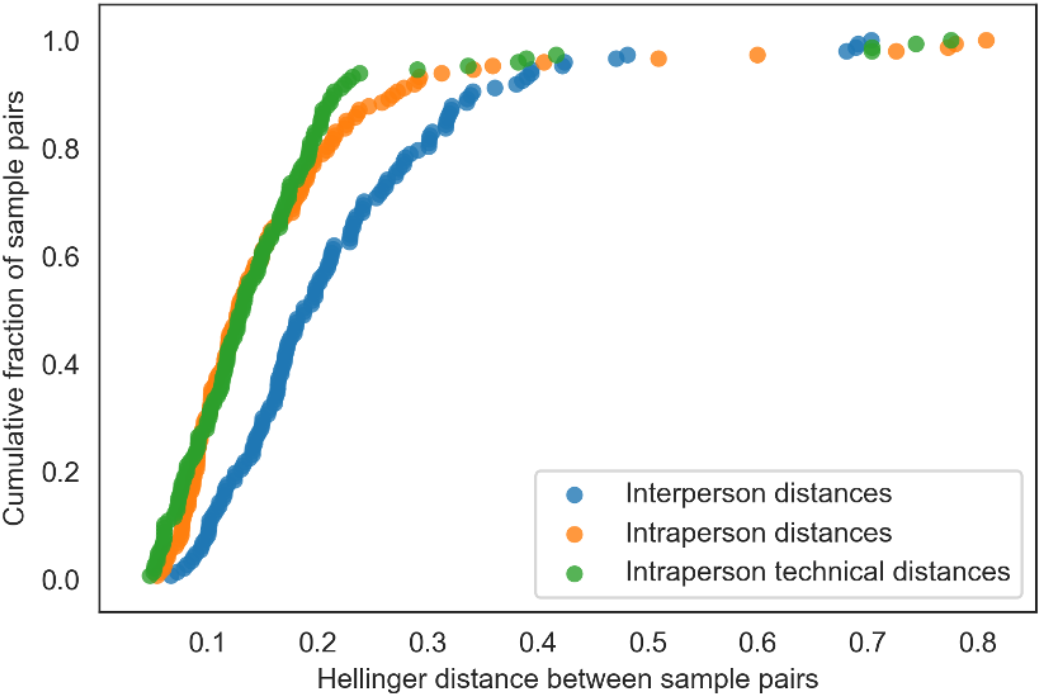
Empirical cumulative distribution function (ECDF) plots of Hellinger distance shows that when taking all genes into account, pairs of samples taken from an individual within one week (green) are not obviously more similar to each other than are samples from one individual taken on two different weeks (orange). However, pairs of samples coming from two different participants (blue) tend to be less similar.

### Most variable genes present in all samples

Using the described test, we analyzed capillary blood from 557 unique individuals. Of all the genes whose transcripts were detected in 100% of the samples, the following genes had the highest variability in gene expression across all samples (Table1). The ability to detect significant differences in expression not only across some of the seldom-detected genes, but also among the most commonly expressed ones, allows for powerful and precise derivation of functional insights specific to an individual’s biology. Gene expression patterns in individuals serve as molecular data input, which inform pathway analysis that yields numerical assessments of levels of activity of specific signaling or metabolic pathway mechanisms of relevance to health and disease. Taken integratively with gut microbial gene expression, complex functional profiles of cross-omic cross-organism molecular pathway interactions offer actionable health insights, such as unique inflammatory, stress response, mitochondrial biogenesis, hormone balance, cardiometabolic, and cellular aging-related themes.

**Table 1.** Most differentially expressed genes detected in all samples. The list of genes with 100% prevalence and coefficient of variability (CV) greater than 1 across all samples is ranked from highest (top) to lowest (bottom) difference in expression across the dataset.

| Ensemble ID | Gene Symbol | Ensemble ID | Gene Symbol |
| --- | --- | --- | --- |
| ENSG00000066739 | ATG2B | ENSG00000072121 | ZFYVE26 |
| ENSG00000070614 | NDST1 | ENSG00000181722 | ZBTB20 |
| ENSG00000105849 | TWISTNB | ENSG00000169118 | CSNK1G1 |
| ENSG00000198887 | SMC5 | ENSG00000152213 | ARL11 |
| ENSG00000025770 | NCAPH2 | ENSG00000076108 | BAZ2A |
| ENSG00000085719 | CPNE3 | ENSG00000118689 | FOXO3 |
| ENSG00000134352 | IL6ST | ENSG00000130764 | LRRC47 |
| ENSG00000181192 | DHTKD1 | ENSG00000130856 | ZNF236 |
| ENSG00000126870 | WDR60 | ENSG00000128829 | EIF2AK4 |
| ENSG00000198265 | HELZ | ENSG00000157601 | MX1 |
| ENSG00000136874 | STX17 | ENSG00000073614 | KDM5A |
| ENSG00000011454 | RABGAP1 | ENSG00000119917 | IFIT3 |
| ENSG00000135503 | ACVR1B | ENSG00000156650 | KAT6B |
| ENSG00000089775 | ZBTB25 | ENSG00000048471 | SNX29 |
| ENSG00000169925 | BRD3 | ENSG00000099290 | WASHC2A |
| ENSG00000114126 | TFDP2 | ENSG00000104219 | ZDHHC2 |
| ENSG00000223609 | HBD | ENSG00000005810 | MYCBP2 |
| ENSG00000141867 | BRD4 | ENSG00000118900 | UBN1 |
| ENSG00000075420 | FNDC3B | ENSG00000132466 | ANKRD17 |
| ENSG00000186174 | BCL9L | ENSG00000176715 | ACSF3 |
| ENSG00000127946 | HIP1 | ENSG00000124177 | CHD6 |
| ENSG00000185129 | PURA | ENSG00000079313 | REXO1 |
| ENSG00000157837 | SPPL3 | ENSG00000132465 | JCHAIN |
| ENSG00000144161 | ZC3H8 | ENSG00000060237 | WNK1 |
| ENSG00000198911 | SREBF2 | ENSG00000131584 | ACAP3 |
| ENSG00000090861 | AARS1 | ENSG00000163214 | DHX57 |

For example, SREBF2 and FOXO3A above vary by as many as 14 units of standard deviation across the dataset, while expressed in all samples, and both play a role in Sirtuin 6 function, which plays a key role in pathways related to positive antioxidant, anti-aging, and metabolic effects (Elhanati et al., 2013; Li et al., 2011; Simeoni et al., 2013; Tao et al., 2013; Xiong et al., 2013). Sirtuin 6 can be one of the intervention strategies for metabolic or aging-related conditions, as it is a known target of many nutrients and herbs. Resveratrol, NAD+ precursors, and fisetin (found in strawberries) are some of the available supplements that can improve Sirtuin function, oxidative stress levels, cellular and mitochondrial health, and have many clinically demonstrated benefits (Oliveira et al., 2017; Wang et al., 2018).

## Discussion

Chronic, non-communicable, diseases are the leading cause of death globally (CDC, 2020). There is strong evidence that human health and chronic diseases, including aging, are heavily influenced by nutrition and microbiome functions. To understand the root causes of chronic diseases, we need to study the human body as an ecosystem, which requires a systems biology approach. Gene expression levels, for both the human and microbiome, play a very important role in chronic diseases onset and progression, and need to be studied in the context of an integrated systems biology platform. While a scalable and clinically validated gut microbiome gene expression test has been developed (Hatch et al., 2019), a human blood transcriptome test with the same features and performance is also needed.

Here we describe a novel whole blood transcriptome test that is clinically validated and globally scalable. The blood sample (50 microliters) can be collected by almost anyone, anywhere, as it uses blood from a finger prick. The samples can be stored and shipped at ambient temperature to a reference laboratory. The method is automated, inexpensive, and high throughput. RPB inactivates pathogens, which is very important for international shipping, as it prevents disease spread. RPB also preserves RNA for up to 28 days and ambient temperature, which obviates the need for cold storage and shipping, which are expensive and not easily available.

Since the majority of published blood transcriptome studies rely on peripheral (venous) blood, we compared the transcriptome data obtained from peripheral and capillary blood and show high gene expression level concordance between the two sample types. This is important in case the results obtained from one sample type need to be compared to the other. We also show that the precision of the test is sufficient to distinguish intra-person technical replicates and longitudinal transcriptome data from other people’s blood transcriptomes.

This test, along with the previously published stool metatranscriptome test and upcoming saliva and vaginal metatranscriptome tests, can be integrated into a comprehensive systems biology platform that can be applied to population-scale longitudinal studies. Such large studies will identify mechanisms of chronic disease onset and progression, improve or bring new diagnostic and companion diagnostic tools, and enable precision nutrition (including probiotics) and microbiome engineering to prevent and cure chronic diseases.

