## Supplemental Figure 1 for "A clinically validated human capillary blood transcriptome test for global systems biology studies"

Supplement

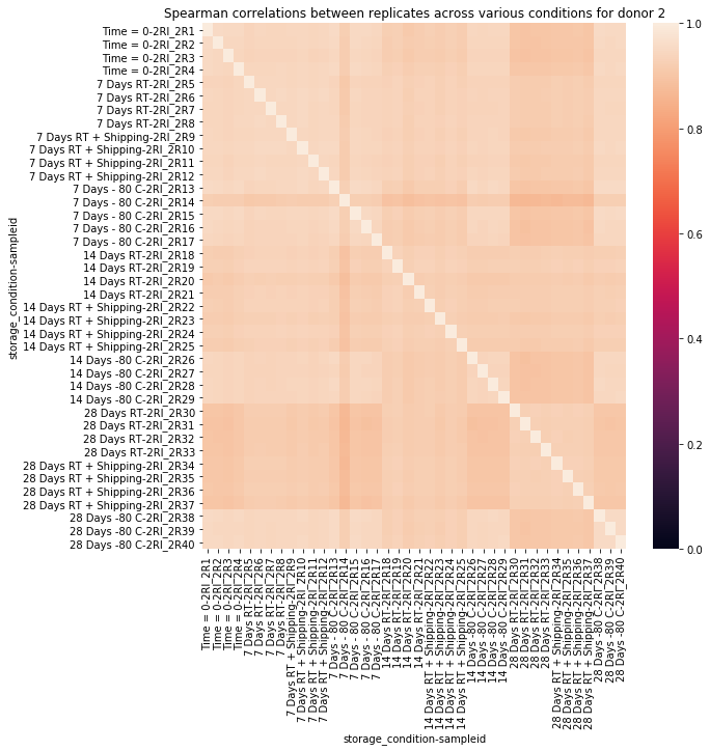

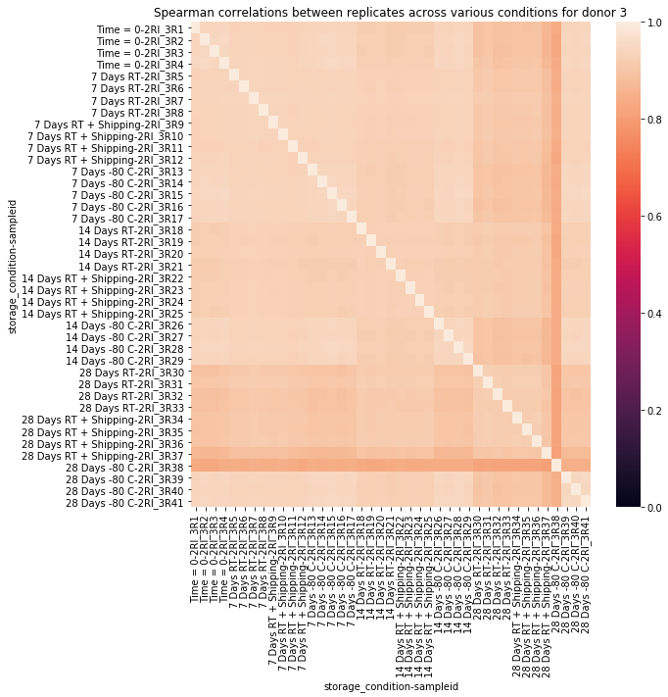

**Figure S1.** Heatmaps showing Spearman correlation coefficients for sample stability of blood samples stored at ambient temperature and -80^o^C for 0, 7, 14, and 28 days with and without shipping. The Spearman correlation coefficients for all conditions in all donor samples were greater than 0.81, showing high concordance. Heatmaps for data obtained from donors 2 and 3 are shown here.
